# Single-Cell Transcriptomic Analysis of the Immune Response to CHIKV Infection

**DOI:** 10.64898/2026.06.24.734421

**Authors:** Peng Huang, Huiyi Wang, Siya Xu, Miaogen Li, Mingjuan Guo, Haiyan Wang, Xiaoyan Gou, Can Wang, Yanjun He, Wei Pan

## Abstract

**Background:** The Chikungunya virus (CHIKV), a re-emerging mosquito-borne alphavirus, is responsible for acute febrile illness and severe polyarthralgia. Although both innate and adaptive immune responses influence the disease outcomes, the detailed cellular immunopathogenesis of CHIKV in peripheral blood is not yet fully elucidated.

**Methods:** We conducted single-cell RNA sequencing (scRNA-seq) on peripheral blood mononuclear cells (PBMCs) obtained from patients acutely infected with CHIKV and from healthy control subjects. Cellular interactions were inferred, and the transcriptomic results were orthogonally validated through quantitative real-time PCR (qPCR) and Enzyme-Linked Immunosorbent Assay (ELISA) to assess systemic interferon-stimulated responses. Furthermore, a comparative analysis was performed using publicly available single-cell data from Dengue virus (DENV) infections.

**Results:** CHIKV infection significantly altered the immune system, increasing monocytes and dendritic cells while reducing T and B lymphocytes. Monocytes and NK cells showed strong activation of interferon-stimulated genes (ISGs). Monocytes were identified as key in driving inflammatory and immune responses. In adaptive immunity, CHIKV led B cells to become plasmablasts with antiviral immunoglobulins and caused T cells and NK-like T cells to show signs of cytotoxicity and exhaustion. Validation showed increased levels of IFN-γ, IFN-β1, MX1, and ISG15. CHIKV triggered a more intense, monocyte-driven interferon response than DENV.

**Conclusions:** Acute CHIKV infection induces a systemic interferon response predominantly centered on monocytes, accompanied by significant alterations in adaptive immunity. Circulating ISG products, including MX1 and ISG15, reflect the transcriptomic activation and may serve as potential biomarkers for assessing the early intensity of innate antiviral responses.

## 1. Introduction

The Chikungunya virus (CHIKV), a re-emerging mosquito-borne alphavirus, poses a significant and increasing threat to global public health, having caused millions of infections worldwide^1–3^. The clinical presentation of the infection typically involves an acute febrile illness, often accompanied by rash, severe polyarthralgia, and myalgia^4^. Notably, approximately 50% of affected individuals develop chronic arthritis, which can persist for several months^5^.

Host immune responses are critical in determining the outcome of CHIKV infection^6^. An immediate innate immune response, particularly the type I interferon (IFN) response, is essential for controlling acute viral replication^7^. However, CHIKV has evolved mechanisms to evade host immune defenses; notably, the viral nonstructural protein 2 (nsP2) disrupts STAT1 signaling and inhibits host transcriptional processes^8,9^. When viral replication exceeds immune control, excessive inflammation may occur, potentially contributing to severe disease manifestations. Unregulated inflammation has been linked to adverse outcomes in CHIKV infection^6^. These findings highlight the importance of elucidating the mechanisms by which host immunity detects and responds to CHIKV infection.

Recent studies emphasize the need for detailed immune profiling in viral infections. In COVID-19, severe cases are linked to myeloid cell issues^10^, while dengue virus (DENV) has distinct immune markers for severe disease^11^. These findings highlight the importance of single-cell analysis in understanding viral immunopathology. However, the immune response in CHIKV remains poorly understood, with research mainly limited to cytokine data^12–14^. Although monocytes^6,15^ and macrophages^16,17^ are involved in CHIKV inflammation, their exact roles are unclear, necessitating a thorough analysis of the immune response in CHIKV.

In this study, we performed single-cell RNA sequencing (scRNA-seq) on peripheral blood mononuclear cells (PBMCs) obtained from patients with acute CHIKV infection to elucidate immune responses at the cellular level. Our analysis identifies a monocyte-centric inflammatory signaling network as a defining characteristic of the CHIKV immune response and reveals significant remodeling within both the innate and adaptive immune compartments. Comparative analysis with DENV infection highlights both shared antiviral responses and virus-specific immune features. Collectively, these findings provide novel insights into the cellular mechanisms underlying CHIKV immunopathogenesis.

## 2. Results

### 2.1 Overall analysis of single-cell PBMC transcriptomic profiling for CHIKV infected cases

To elucidate the immunological landscape of PBMCs in individuals infected with CHIKV, we conducted scRNA-seq utilizing the 10x Genomics platform on PBMC samples obtained from 10 healthy donors and 29 CHIKV-infected patients (Fig. 1A). Rigorous quality control measures were applied to the single-cell dataset to exclude low-quality cells and genes with low expression levels, resulting in the retention of 414,257 cells and 35,168 genes for subsequent analyses. Through unsupervised clustering, we delineated 10 principal immune and hematopoietic cell populations, characterized by the expression of canonical marker genes. These populations comprised CD4⁺ T cells, CD8⁺ T cells, B cells, plasma cells, natural killer (NK) cells, γδ T cells, dendritic cells (DCs), monocytes, erythroid cells, and platelets (Fig. 1B-C). The distribution of cells across samples was relatively balanced, thereby minimizing potential biases in downstream analyses (Fig. 1D).

**Figure 1.**
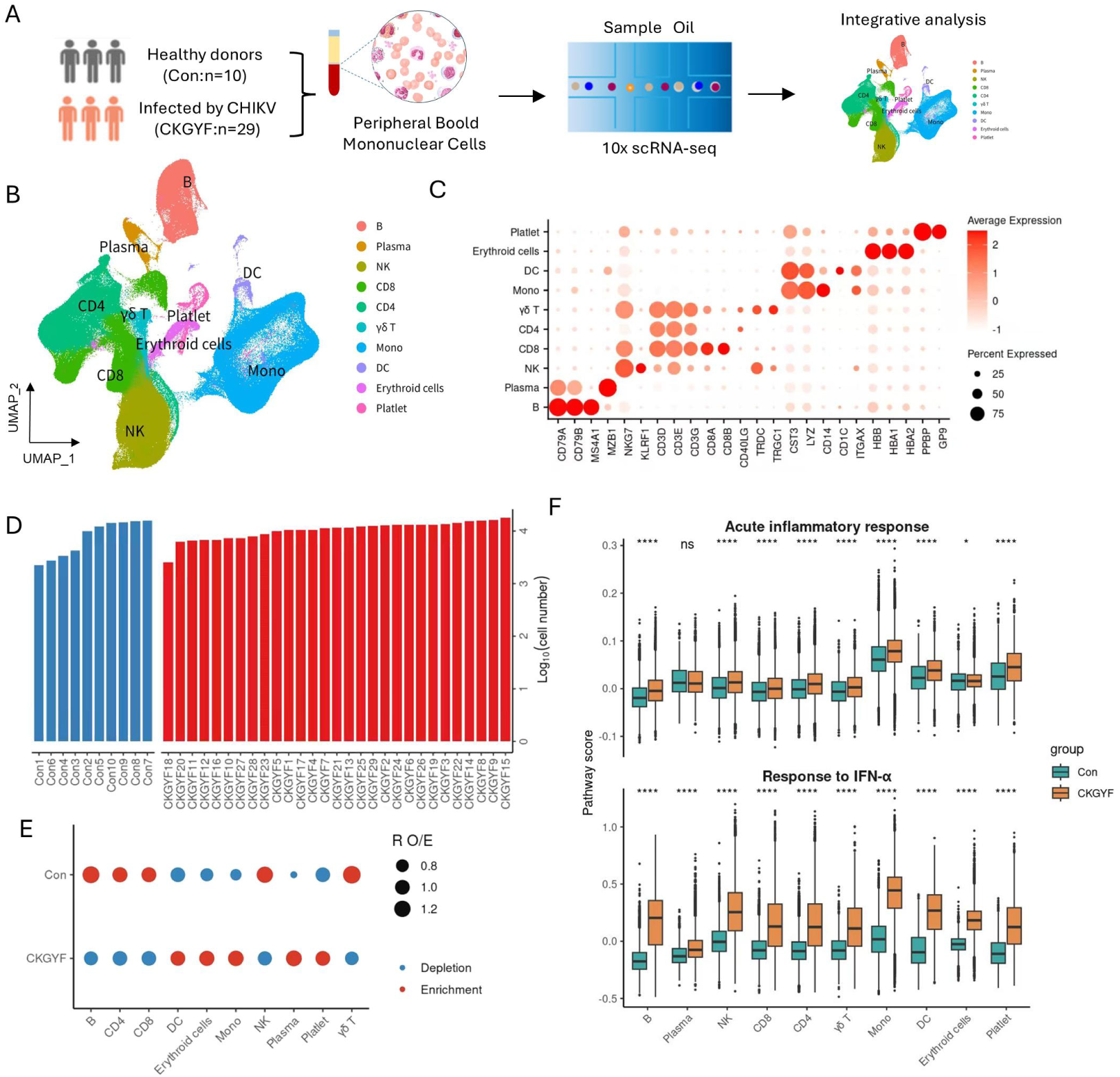
Overview of the study design and single-cell transcriptome profiling of peripheral immune cells.(A) Study overview schematic, including data from 10 healthy controls and 29 CHIKV-infected patients. **(B)** UMAP clustering of single-cell transcriptome data from CHIKV-infected samples. **(C)** Typical marker genes used for cell type annotation in the UMAP clustering. **(D)** Bar plot showing the number of cells profiled from each individual sample. **(E)** Observed/Expected (O/E) ratio plot illustrating the enrichment preference of major cell types between groups. **(F)** Group differences in acute inflammatory and interferon response pathway scores of major cell types.

Subsequently, we evaluated alterations in immune cell composition between patients infected with CHIKV and healthy control subjects by employing the ratio of observed to expected (O/E) cell counts. In comparison to healthy donors, individuals with CHIKV infection demonstrated a significant enrichment of DCs, monocytes, erythroid cells, plasma cells, and platelets, while B cells, CD4⁺ T cells, and CD8⁺ T cells were relatively depleted (Fig. 1E). To explore the antiviral and pathogenic immune responses associated with CHIKV infection, we analyzed the expression levels of two critical pathways, specifically the Gene Ontology Biological Process (GOBP) terms: response to interferon (IFN)-α and acute inflammatory response, across major cell types under both conditions. Our findings revealed that the response to IFN-α was consistently and significantly upregulated in all major cell types derived from the PBMCs of CHIKV-infected patients (Fig. 1F). Furthermore, with the exception of plasma cells, the acute inflammatory response was significantly upregulated across all cell types under both conditions (Fig. 1F).

### 2.2 Strong interferon responses were observed in innate immune cells

To elucidate the regulatory effects of CHIKV infection on host innate immune cells, monocytes and DCs were further stratified into distinct subpopulations. Our analysis identified multiple transcriptionally distinct subsets of innate immune cells, including NK cells, CD1C⁺ conventional dendritic cells (cDC2), plasmacytoid dendritic cells (pDCs), CD14⁺ classical monocytes (Mono_CD14), CD16⁺ non-classical monocytes (Mono_CD16), and NEAT1-high monocytes (Mono_NEAT1) (Fig. 2A-B). We evaluated the subset-specific enrichment preferences of innate immune cells in response to CHIKV infection. Compared to healthy controls, patients infected with CHIKV exhibited a modest enrichment of pDCs, Mono_CD14, Mono_CD16, and Mono_NEAT1 cells, along with a decreased abundance of cDC2 cells (Fig. 2C-D). Notably, the inflammatory scores of all innate immune cell populations were elevated in the CHIKV-infected cohort, with Mono_CD14 and Mono_CD16 demonstrating the highest levels. Furthermore, other PBMC populations also showed a general trend of increased inflammatory activity (Fig. 2E). Collectively, these findings suggest that CHIKV infection induces a broadly heightened inflammatory state within the innate immune compartments.

**Figure 2.**
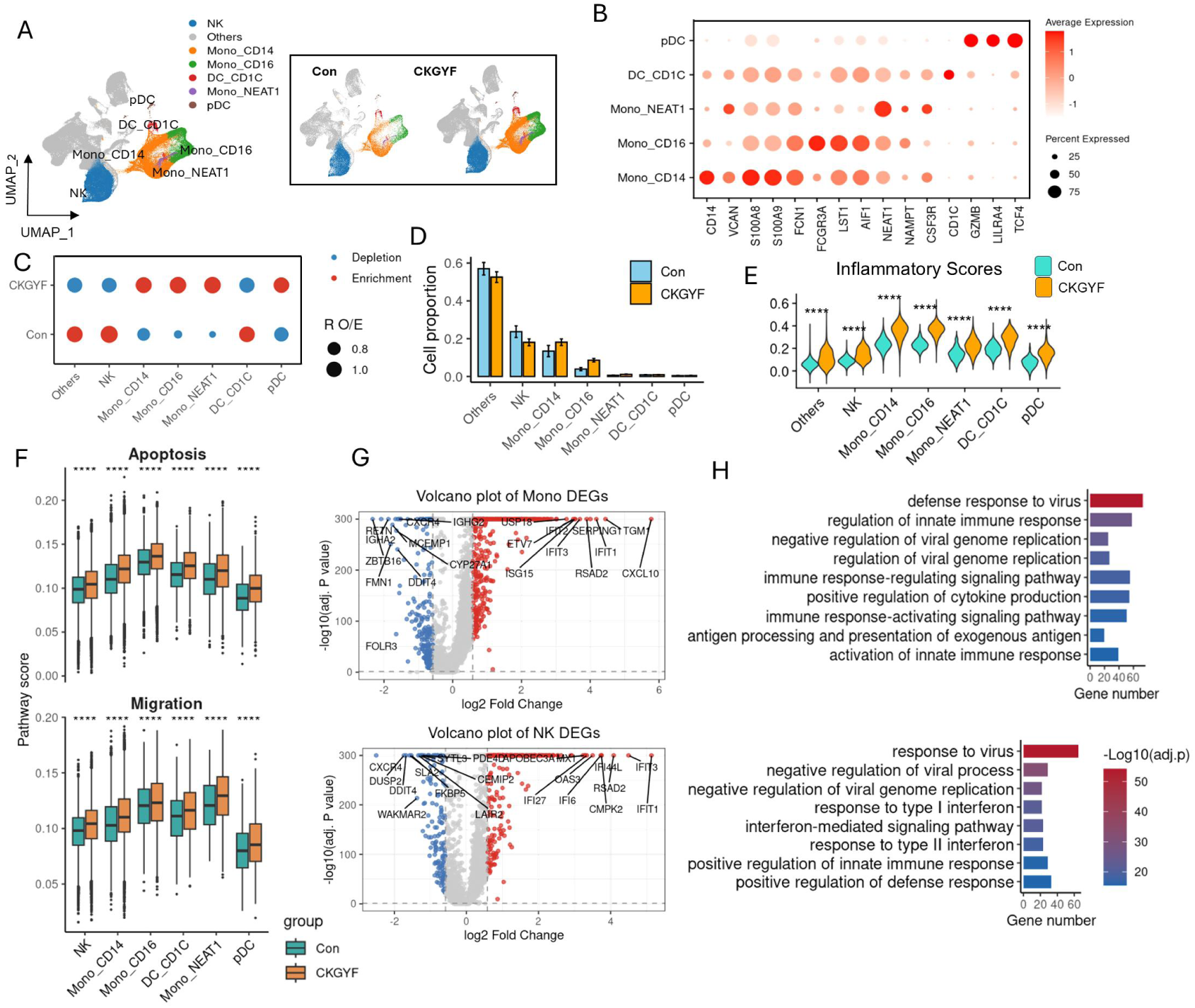
Single-cell transcriptomic profiling and functional characterization of innate immune cells. **(A)** UMAP projection of innate immune cell subsets. Each dot corresponds to a single cell, colored according to cell type. **(B)** Dot plot showing the expression of typical marker genes for the annotation of each innate immune cell subset. **(C)** Observed/Expected (O/E) ratio plot illustrating the enrichment preference of major innate immune cell types between groups. **(D)** Group changes in the proportion of innate immune cell types. **(E)** Group differences in inflammatory signature enrichment scores of major innate immune cell types. **(F)** Volcano plot of differentially expressed genes (DEGs) in monocytes and NK cells between groups. **(G)** GOBP enrichment analysis for the identified DEGs. **(H)** Comparing the group differences in enrichment scores of apoptosis-related pathway and migration-related pathway for major innate immune cell types.

To further elucidate the transcriptomic alterations in innate immune cells following CHIKV infection, we conducted a comparative analysis of gene expression profiles between infected and healthy conditions in monocytes and NK cells. Our findings demonstrated a significant upregulation of interferon-stimulated genes (ISGs), such as IFIT1 and IFIT3, in both monocytes and NK cells. GOBP enrichment analysis indicated that the DEGs in these cell types were significantly enriched in pathways related to the regulation of innate immune responses and antiviral defense mechanisms. Additionally, scores for migration and apoptosis were elevated in innate immune cells from the infected cohort (Fig 2F-H).

### 2.3 Immunological features of B cell subsets

In order to elucidate the dynamic alterations among various B cell subtypes, we identified six transcriptionally distinct B cell subsets: naive B cells (B_Naive), germinal center B cells (B_GC), memory B cells (B_Memory), intermediate memory B cells (B_iMemory), plasmablasts (B_Plasmablast), and plasma cells (B_Plasma) (Fig. 3A-B). Comparative analysis of their relative proportions demonstrated a significant increase in B_Plasmablast and B_Plasma populations within the infected cohort, accompanied by a notable decrease in B_Naive cells. This suggests that CHIKV infection facilitates the functional transition of B_Naive cells towards B_Plasmablast (Fig. 3C). To further explore transcriptomic alterations, we analyzed the expression profiles of B/plasma cells under infected conditions relative to healthy controls (HD). Several classical ISGs, such as *IFIT1*, *IFIT3*, *IFI44L*, *RSAD2*, *CMPK2*, and *CXCL10*, were significantly upregulated (Fig. 3D). Additionally, DEGs most significantly enriched in CHIKV-infected patients were associated with defense responses to viral or symbiotic challenges (Fig. 3E). These observations underscore the robust activation of interferon-mediated antiviral responses in B cells following CHIKV infection.

**Figure 3.**
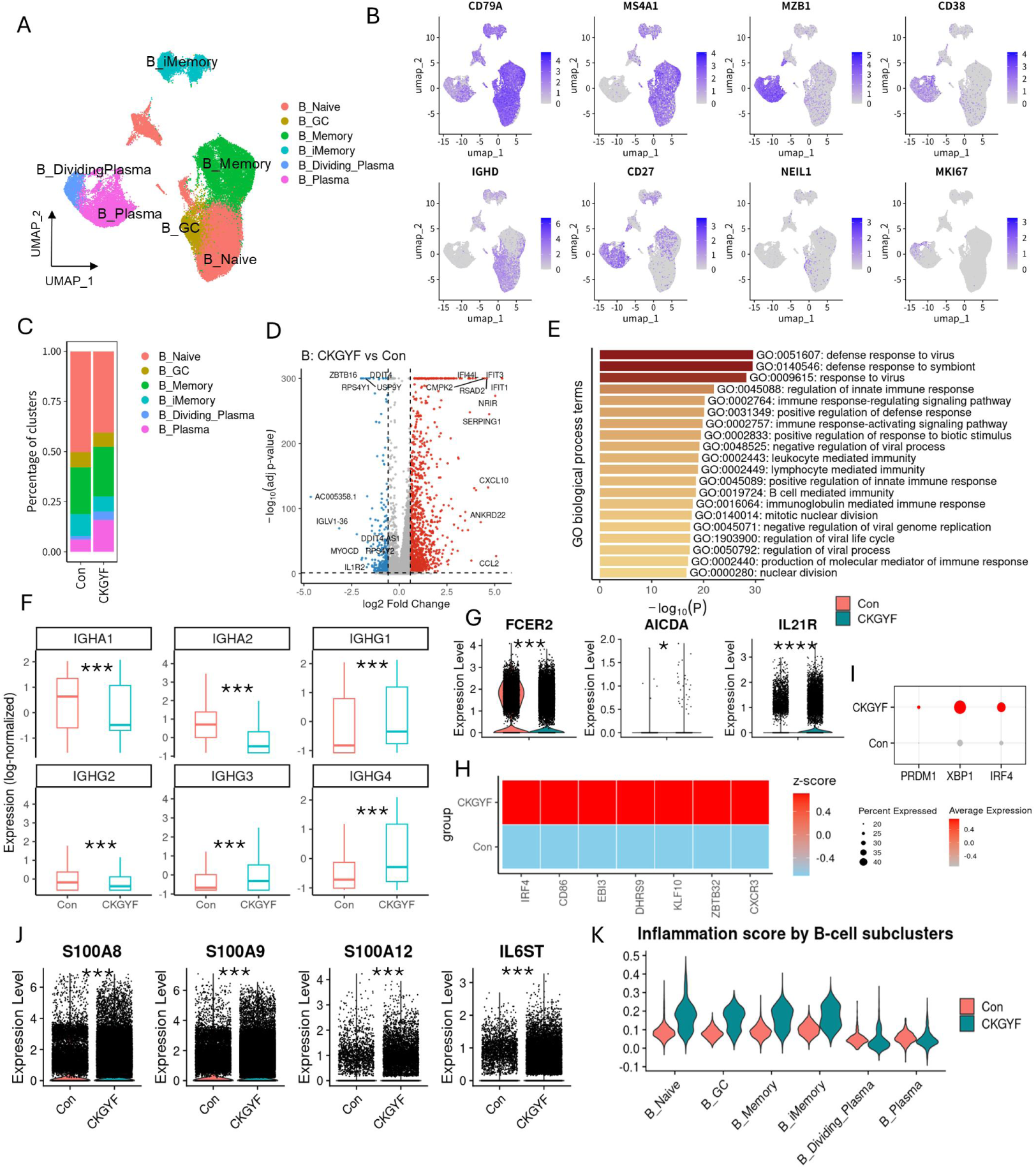
Immunological features of B cell subsets. **(A)** UMAP projection of B cell subsets. **(B)** Typical marker genes used for the annotation of B cell subsets. **(C)** Relative proportion of each B cell subset between groups. **(D)** Differentially expressed genes (DEGs) in B cells between the CHIKV-infected group and healthy control group. **(E)** GOBP enrichment analysis of upregulated DEGs. **(F)** Group differences in antibody secretion profiles of plasma cells. **(G)** Expression changes of activation-related genes in B_Naive. **(H)** Expression changes of activation-related genes in B_Memory. (I) Expression changes of genes related to antibody synthesis and secretion in plasma cells from the CHIKV-infected group compared with the healthy control group. **(J)** Expression changes of multiple cytokines in B cells from the CHIKV-infected group compared with the healthy control group. **(K)** Differences in inflammatory scores of B cell subsets between the CHIKV-infected group and the healthy control group.

The gene expression patterns associated with antibody secretion demonstrated notable alterations in immunoglobulin isotype profiles subsequent to CHIKV infection. Specifically, there was a significant upregulation of *IGHG1*, *IGHG3*, and *IGHG4*, while *IGHA1*, *IGHA2*, and *IGHG2* were downregulated in the CHIKV-infected cohort compared to healthy controls (Fig. 3F). Collectively, these findings imply a shift in antibody production within plasma cells towards an immunoglobulin isotype profile that is more oriented towards antiviral activity following CHIKV infection. Furthermore, the expression levels of critical activation genes, *AICDA* and *IL21R*, were markedly upregulated in naive B cells, while the core transcription factors *PRDM1*, *XBP1*, and *IRF4*, which are associated with plasma cell differentiation, were significantly elevated in the CHIKV-infected group (Fig. 3G, 3I). In conclusion, the expression levels of genes associated with inflammation, such as *S100A8*, *S100A9*, *S100A12*, and *IL6ST*, were markedly upregulated in B cells, indicating a pronounced inflammatory activation state following infection (Fig. 3J, 3K).

### 2.4 Features of T cell subsets in patients with CHIKV

To elucidate alterations in individual T cell subsets across different conditions, we performed subclustering of T cells derived from PBMCs, resulting in the identification of 15 distinct subsets based on the expression and distribution of canonical T cell markers (Fig. 4A-B, Fig.S1, Fig.S2). These subsets comprised 8 subtypes of CD4^+^ T cells, 4 subtypes of CD8^+^ T cells, and 3 subtypes of NK-like T cells characterized by the markers CD3E^+^, CD40LG^-^, CD8A^-^, and TYROBP^+^. We conducted a comparative analysis of T cell subset distribution between the control group and the CKGYF group, revealing a significant enrichment of CD56^+^ NK-like T cells (NK-like T_CD56) in the CKGYF group (Fig. 4C). Furthermore, we assessed the cytotoxicity and exhaustion scores of effector subsets. The CKGYF group exhibited significantly elevated cytotoxicity and exhaustion scores compared to the control group (Fig. 4D), suggesting an enhancement of antiviral effector activity coupled with progressive T cell exhaustion in response to CHIKV infection. Following CHIKV infection, NK-like T_CD56 cells demonstrated a reduction in the expression of naïve T cell markers, alongside an upregulation of genes associated with cytotoxicity and exhaustion. GOBP enrichment analysis of NK-like T_CD56 marker genes revealed significant enrichment in pathways related to leukocyte-mediated cytotoxicity and cell activation (Fig 4E-F). Pathway enrichment analysis of upregulated DEGs in T cells indicated the most notable enrichment in pathways involved in defense response to viruses, positive regulation of cytokine production, and response to interferon (Fig 4G-H). Furthermore, both global and subset-level analyses revealed significantly elevated apoptosis and migration scores in T/NK-like T cells within the CKGYF group, suggesting enhanced trafficking towards inflammatory sites and activation-induced cell death during the contraction phase of the antiviral immune response (Fig 4I-J).

**Figure 4.**
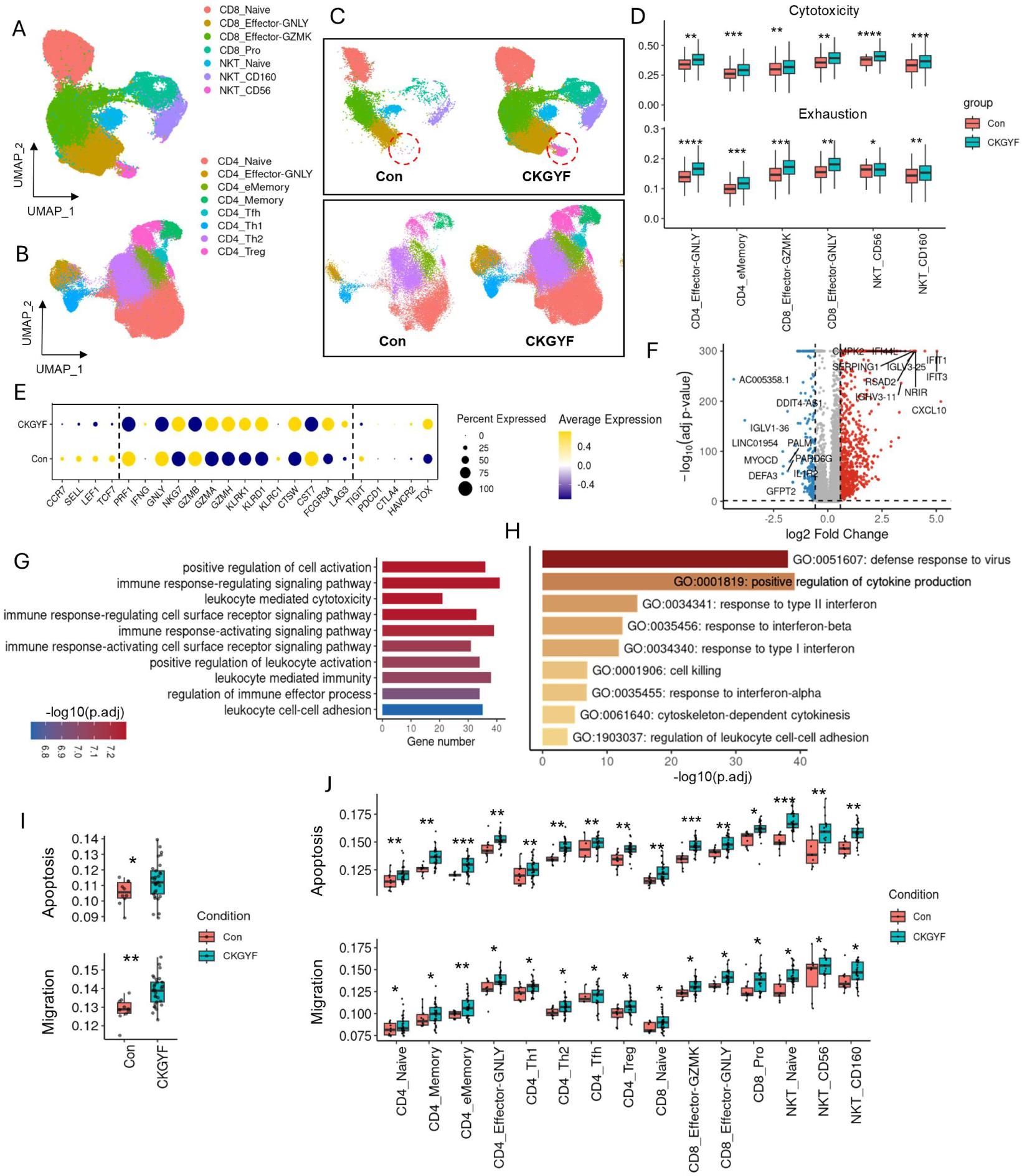
Single-cell transcriptomic profiling and functional characterization of T cell and NKT cell subsets during CHIKV infection. **(A)** UMAP projection of CD8⁺T/NKT cell subsets. **(B)** UMAP projection of CD4⁺T cell subsets. **(C)** UMAP plot showing the group differences in T cell distribution between the healthy control and CHIKV-infected groups. **(D)** Differential analysis of cytotoxicity and exhaustion scores for effector-related T cell subsets. **(E)** Expression profile of cytotoxicity- and exhaustion-related differentially expressed genes in the NKT_CD56 subset. **(F)** Functional enrichment analysis of marker genes for the NKT_CD56 subset. **(G)** Volcano plot of differentially expressed genes (DEGs) in T cells between groups. **(H)** Pathway enrichment analysis of upregulated genes in T cells. **(I)** Group differences in apoptosis and migration scores of total T cells. **(J)** Group differences in apoptosis and migration scores of each T cell subset.

### 2.5 CHIKV infection enhances monocyte-centered immune cell communication networks

To investigate the potential intercellular relationships and transcriptomic alterations, the differences in cellular interactions among cells in CHIKV-infected and uninfected samples were analyzed using CellChat. Following infection, monocytes emerged as the central signaling hub, demonstrating significantly increased incoming and outgoing communication strength, as well as a higher frequency of communication events, particularly with B cells, plasma cells, NK cells, and γδ T cells (Fig 5A, 5B). In the CKGYF group, TNF signaling from monocytes to CD8^+^ T and B cells was notably enhanced, and the SIGLEC1-SPN ligand-receptor interaction from monocytes to other immune cells was also intensified, potentially facilitating immune cell recruitment and activation during infection (Fig 5C). Comparative analysis of monocyte-mediated signaling pathways revealed a significant upregulation of pathways related to inflammation, immune recruitment, and immune regulation (such as CXCL, BAFF, TNF, IL, ICAM/SELPLG), underscoring that CHIKV infection induces heightened monocyte-driven inflammatory and immunoregulatory signaling (Fig 5D). We conducted an in-depth analysis of the principal communication pathways originating from monocytes and discovered that pathways related to MHC-I and MHC-II were predominantly influenced by interactions between various HLA-CD8 and HLA-CD4 molecules. This finding suggests that antigen presentation-mediated T cell activation constitutes a predominant mechanism of cel-cell communication under pathological conditions (Fig 5E).

**Figure 5.**
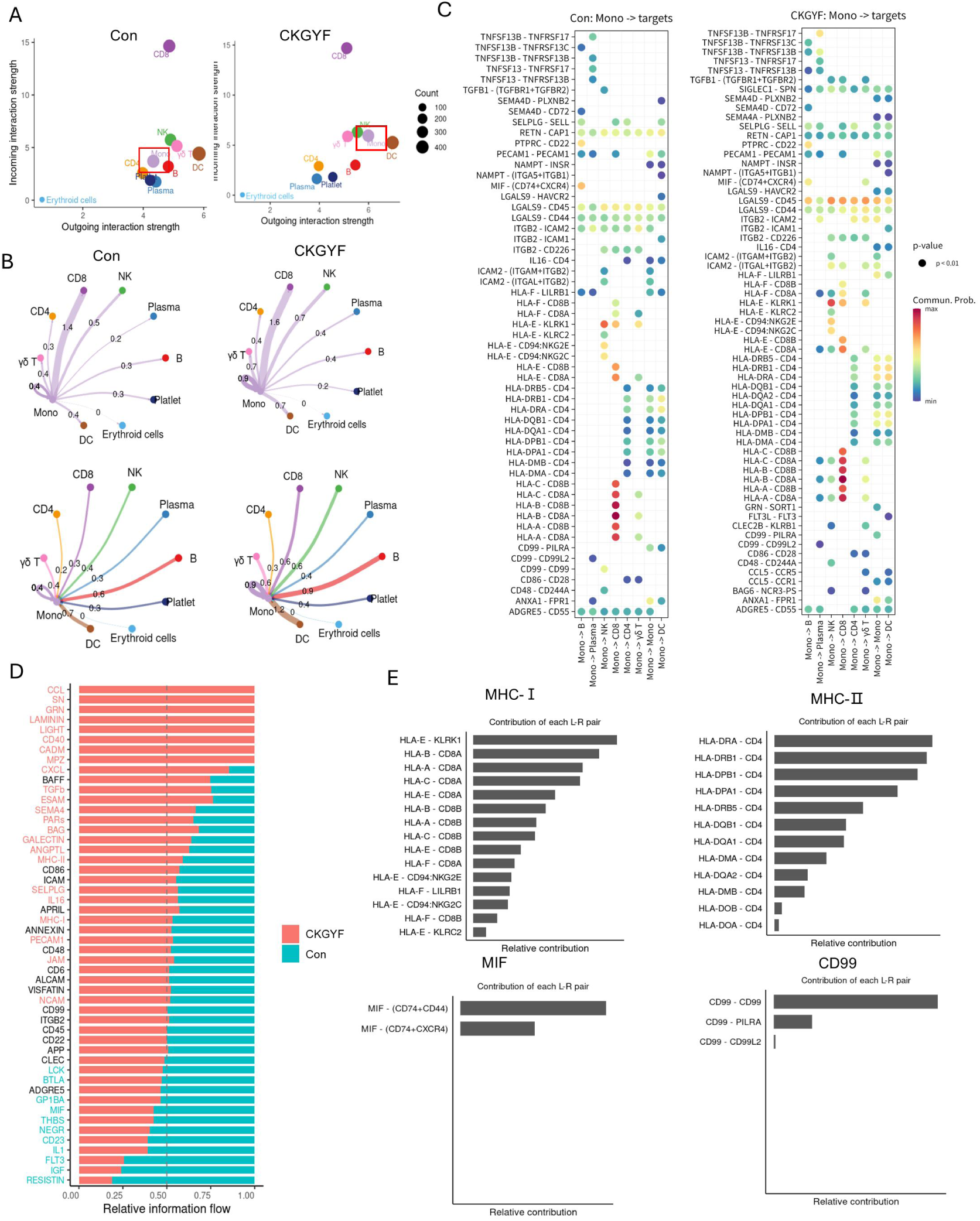
CHIKV infection enhances monocyte-centered immune cell communication networks. **(A)** Scatter plot showing outgoing and incoming interaction strength of immune cell subsets in the CKGYF and uninfected groups. **(B)** Differential interaction strength between CKGYF and uninfected groups. **(C)** Dot plot showing LR pairs interactions transmitted from monocytes to other cell types in the CKGYF and uninfected groups. **(D)** Comparison of outgoing signaling pathways from monocytes between the CKGYF and uninfected groups. **(E)** Dissection of the major monocyte-derived communication signaling pathways in the CKGYF group was performed to evaluate the relative contribution of individual LR pairs to the overall pathway strength.

### 2.6 Distinct antiviral immune responses and systemic validation of the interferon-driven state

Subsequent analyses revealed that IgM was the predominant isotype in the host antibody response following CHIKV infection, with IGHM constituting 78.72% of the total immunoglobulin expression (Fig. 6A). We then evaluated interferon production across major PBMC types. Both the proportion of *IFNG*-positive cells and the mean expression level of *IFNG* were significantly elevated in CD8^+^ T cells and NK cells compared to other immune cell subsets (Fig. 6B-C). To systematically compare immune responses between CHIKV and DENV infections, we integrated our CHIKV single-cell dataset with a publicly available DENV dataset (GSE220969) (Fig. 6D). Based on the ISG expression score, monocytes were identified as the primary contributors to the differential ISG expression between the two groups. Specifically, monocytes in the CHIKV group exhibited significantly higher ISG expression levels than those in the DENV group (Fig. 6E-F). Whole-transcriptome differential analysis further confirmed substantial divergence in the transcriptomic profiles between the CHIKV and DENV groups (Fig. 6I). GOBP enrichment analysis of the intersecting genes that were upregulated in the CHIKV group, in comparison to both the control and DENV groups, revealed a significant enrichment in antiviral immune pathways (Fig. 6J). Furthermore, monocytes from the CHIKV group exhibited a notably increased expression of genes associated with antigen presentation, including *HLA-DQA1*, *IFI30*, and *UNC93B1*, relative to the DENV group. Additionally, the inflammatory scores of these monocytes were substantially elevated (Fig. 6K-L).

**Figure 6.**
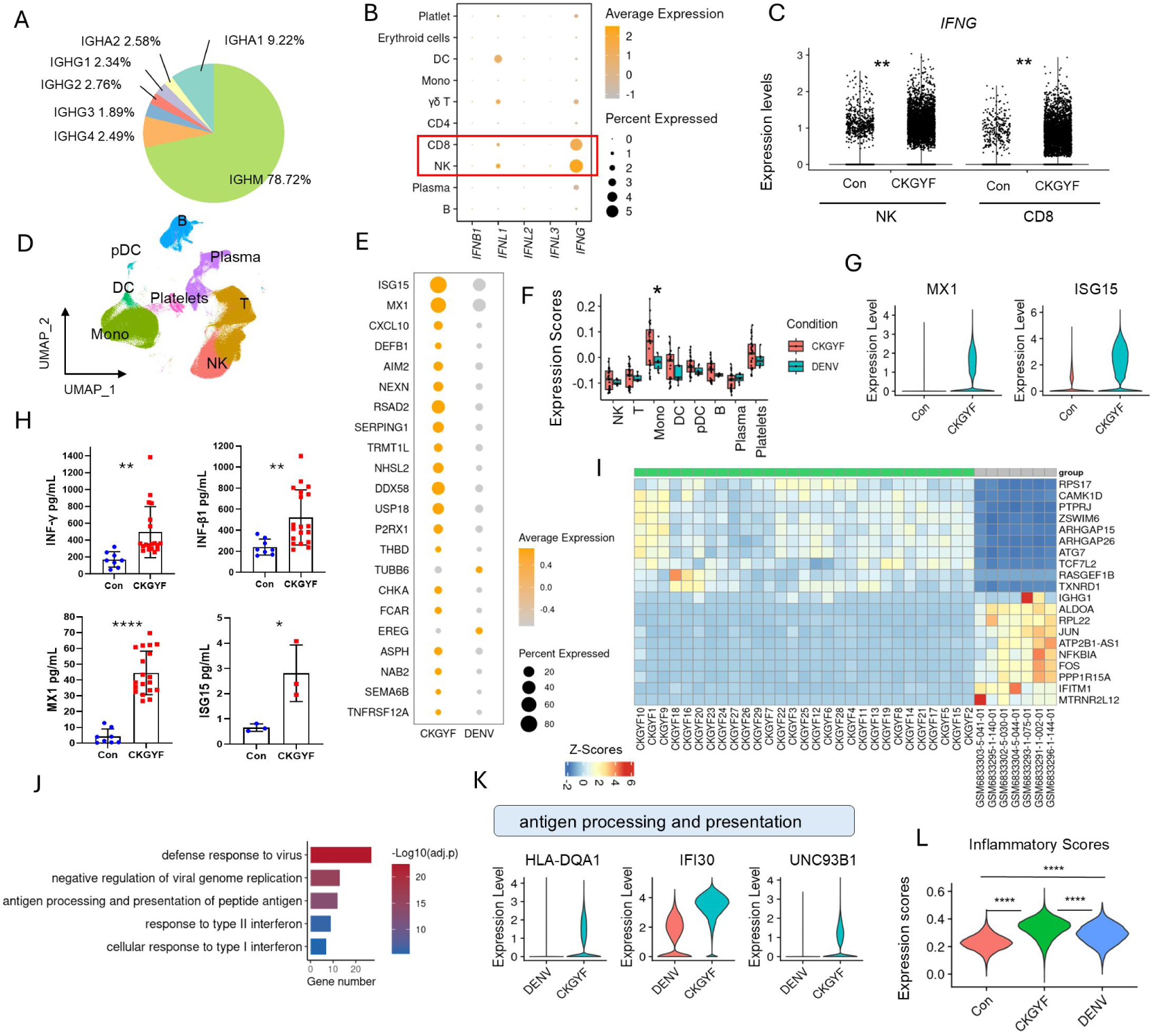
Comparative analysis of host immune response characteristics between CHIKV and DENV infection. **(A)** Proportion of antibody isotype secretion in the CHIKV group. **(B)** Dot plot showing the interferon secretion profile of major cell types. **(C)** Vlnplot showing the group differences in *IFNG* expression in NK cells and CD8⁺ T cells. **(D)** Integrated clustering plot after merging single-cell datasets from the CHIKV group and DENV group (GSE220969). **(E)** Group differences in the expression of interferon-stimulated genes (ISGs). **(F)** Group differences in ISGs expression scores across distinct cell types. **(G)** Differential expression profile of *MX1* and *ISG15* in PBMCs between Con and CKGYF. **(H)** ELISA analysis of IFN-γ, IFN-β1, and MX1 levels in patient plasma samples from the Con (n = 8) and CKGYF (n = 19) groups, and ISG15 levels in a subset of these samples from the Con (n = 3) and CKGYF (n = 3) groups. **(I)** Heatmap of the top 10 upregulated and downregulated differentially expressed genes between the CHIKV and DENV groups. **(J)** GOBP enrichment analysis of the intersecting genes upregulated in the CHIKV group compared with the control and the DENV group were prominently enriched in antiviral immune pathways. **(K)** Differential expression profile of antigen presentation-related genes in monocytes. **(L)** Differences in inflammatory scores of monocytes among the Con, CKGYF and DENV groups.

To independently corroborate the robust interferon-stimulated transcriptomic signatures identified in silico at the systemic level, we evaluated both mRNA expression in bulk PBMCs and circulating protein concentrations in patient plasma. qPCR analysis demonstrated a significant upregulation in the mRNA levels of type I and II interferons (*IFNB1*, *IFNG*) and classical interferon-stimulated genes (*MX1*, *ISG15*) in PBMCs from patients infected with CHIKV compared to healthy controls (Figure 6G). Remarkably, consistent with this transcriptional upsurge, our concurrent ELISA analysis confirmed that the circulating protein concentrations of IFN-β1, IFN-γ, MX1, and ISG15 were significantly elevated in the patient cohort (Figure 6H). Notably, the systemic increase in IFN-γ reflects the enhanced effector functions of CD8^+^ T and NK cells identified in our single-cell atlas, while the pronounced induction of IFN-β1, MX1, and ISG15 supports the widespread antiviral state primarily mediated by the monocyte network. Collectively, this orthogonal validation from transcript to protein provides compelling evidence that acute CHIKV infection triggers a substantial, systemic interferon-driven inflammatory response.

## 3. Discussion

In this study, we conducted a comprehensive characterization of the peripheral immune landscape associated with CHIKV infection at a single-cell resolution, utilizing PBMC samples from both healthy individuals and infected patients. Our findings demonstrate that CHIKV infection significantly alters the circulating immune compartment, characterized by a pronounced enrichment of innate immune populations, particularly monocytes and DCs, alongside a reduction in conventional T- and B-cell populations. Across major cell types, there was a widespread elevation of interferon-related signatures, suggesting that CHIKV infection triggers a systemic antiviral transcriptional program in peripheral blood. Notably, among innate immune cells, monocyte subsets exhibited the most robust inflammatory and interferon-stimulated responses, underscoring the pivotal role of monocytes in orchestrating the host’s antiviral defense during CHIKV infection. Within the B-cell lineage, CHIKV infection was associated with an expansion of plasmablasts and modifications in immunoglobulin isotype profiles, indicative of an antiviral-oriented humoral response. Within the T-cell compartment, we observed heightened cytotoxicity and exhaustion signatures, notably within effector-associated T/NK-like T cell subsets.

These observations collectively imply that CHIKV infection induces interferon-mediated activation of innate immune responses, thereby enhancing antiviral defense strategies. This response pattern parallels the innate immune activation observed in PBMCs during COVID-19 infection^18^. Furthermore, DEGs in monocytes were particularly enriched in pathways associated with the positive regulation of cytokine production, suggesting that activated monocytes may further influence immune responses through cytokine modulation.

Simultaneously, there was a notable upregulation of IGHG1, IGHG3, and IGHG4, while IGHA1, IGHA2, and IGHG2 exhibited downregulation in the cohort infected with CHIKV. Notably, IgG1 and IgG3 are well-recognized for their potent antiviral effector functions and their capacity to activate complement pathways^19,20^, whereas IgG2 is primarily associated with responses to bacterial capsular polysaccharide antigens^21^. Furthermore, IgG4 is characterized by its immunomodulatory properties^22^. Collectively, these observations suggest a shift in antibody production within plasma cells towards an immunoglobulin isotype profile that is more aligned with antiviral activity following CHIKV infection.

The observed elevation in plasma cell numbers, concomitant with a decrease in the overall B-cell population, likely indicates an accelerated differentiation process of B cells into antibody-secreting cells. This mechanism leads to the depletion of the conventional B-cell reservoir and the expansion of plasma cells. Such a phenomenon is frequently documented in arboviral infections, where rapid humoral responses are initiated during the acute phase. For instance, research on dengue virus infection has demonstrated that the infection can induce the expansion of CD14⁺, CD16⁺monocytes, which subsequently promote B-cell differentiation into plasmablasts and plasma cells^23^. Furthermore, the enrichment of platelets and erythroid cells (presumably nucleated erythroid progenitors), along with enhanced antiviral and inflammatory responses, suggests that platelet activation^24^ and stress hematopoiesis^25^ may occur during CHIKV infection. Overall, our findings suggest that CHIKV infection triggers a coordinated immune response characterized by robust innate antiviral activation and dynamic remodeling of the adaptive immune system.

A notable discovery of this study is the identification of monocytes as the central communication hub within the CHIKV immune network. Utilizing CellChat^26^ analysis, it was revealed that CHIKV infection substantially enhances monocyte-centered signaling, with significant amplification of the TNF, CXCL, BAFF, and IL pathways. These findings contribute to the current understanding of CHIKV immunopathogenesis by suggesting that monocytes serve not only as transcriptionally activated responders but also as crucial coordinators of immune communication in the peripheral circulation. Furthermore, our comparative analysis with DENV^11^ underscores the immunological distinctiveness of CHIKV infection. Although both arboviral infections trigger antiviral immune responses^27^, CHIKV is characterized by more pronounced interferon-stimulated gene expression and increased antigen presentation in monocytes.

In addition, our orthogonal validation, which integrates cellular transcriptomics with circulating protein phenotypes, establishes a robust framework for elucidating systemic CHIKV pathology. The remarkable concordance between the upregulated monocyte-driven ISG network identified through scRNA-seq and the systemic elevations of MX1 and ISG15 quantified via qPCR and ELISA is particularly noteworthy. While traditional clinical assessments of CHIKV primarily focus on broad inflammatory markers^28^, our findings underscore the potential of circulating MX1 and ISG15 as precise, mechanistically grounded biomarkers for assessing the intensity of the early innate antiviral response. Furthermore, given the significantly elevated migration scores and enhanced chemotactic signaling, such as the CXCL network^29,30^, observed in these highly inflammatory monocytes, we propose that these peripheral cells function as a “vanguard” population. Upon systemic activation, these circulating inflammatory monocytes may efficiently migrate to local musculoskeletal sites, potentially contributing to the synovial inflammation and severe polyarthralgia that are clinical hallmarks of CHIKV infection^31,32^.

Despite these insights, it is important to acknowledge several limitations inherent in this study. Firstly, the analysis was conducted using PBMC samples, which, while providing a convenient overview of systemic immune responses, do not fully capture the tissue-specific immune events occurring at local sites of viral replication and pathology, such as the joints. Secondly, the study employed a cross-sectional design, which restricts our ability to elucidate the temporal dynamics of immune responses throughout the course of infection and recovery. Thirdly, although multiple transcriptional alterations and inferred changes in cell-cell communication were identified, these developmental inferences remain fundamentally descriptive in the absence of pseudotime trajectory mapping or in vitro functional cellular assays. Lastly, the comparative integration with a public DENV dataset may be subject to analytical batch effects, despite the application of integrative methods. Future research should incorporate longitudinal multi-omics approaches and targeted functional studies to enhance the refinement of pathogen-specific immune biomarkers.

## 4. Materials and methods

### Patient Recruitment and Sample Preparation

This study received approval from the Institutional Ethics Committee of Foshan Women and Children’s Hospital (Approval No.: FSFY-MEC-2025-138). Between July 2025 and September 2025, 29 confirmed cases of Chikungunya fever and 10 healthy control subjects were recruited. The inclusion criteria for patients infected with CHIKV were: (i) the presence of fever accompanied by severe arthralgia; (ii) a positive result for CHIKV RNA via real-time reverse transcription polymerase chain reaction (real-time RT-PCR) or a positive result for serum anti-CHIKV IgM antibody; and (iii) symptom onset within 7 days prior to enrollment. Healthy controls were individuals undergoing routine health examinations during the same timeframe, with no history of CHIKV infection or febrile symptoms. Peripheral venous blood samples (5 mL) were collected from each participant into EDTA-anticoagulated tubes. The samples were allowed to stand at room temperature for 30 minutes, followed by centrifugation at 3000 rpm for 10 minutes at 4°C to separate the plasma from the cellular fraction. The plasma was aliquoted and stored at −80°C for subsequent ELISA analysis, while the cellular fraction was utilized for the isolation of PBMCs.

### scRNA-seq data processing, cell clustering, and dimension reduction

The processing of scRNA-seq data, including cell clustering and dimensionality reduction^33,34^, involved aligning the sequenced reads to the GRCh38 human reference genome using Cell Ranger (version 3.1.0, 10x Genomics). To mitigate potential ambient RNA contamination and random barcode swapping in the raw UMI-based scRNA-seq data, the remove-background function in CellBender was employed. Further quality control of the cells was conducted, retaining those with 200 to 6,000 expressed genes and less than 15% mitochondrial RNA content.

Downstream data integration, cell clustering, and dimensionality reduction were conducted utilizing Seurat (version 5). In summary, 2,000 highly variable genes (HVGs) were selected for subsequent analysis. To mitigate technical batch effects across different samples, canonical correlation analysis (CCA) was employed during the data integration process. Principal component analysis (PCA) was performed using the top 30 components derived from the identified HVGs. Cells were then clustered using a nearest-neighbor graph approach, and cluster-specific marker genes were identified using the MAST algorithm. Finally, the clustered cells were visualized in a two-dimensional space through the application of the non-linear dimensionality reduction technique, UMAP.

### Analysis of Differentially Expressed Genes (DEGs)

DEGs were identified utilizing the MAST algorithm as implemented in the Seurat package^35^ (version 5). Pairwise comparisons were performed across various conditions or cell clusters. Genes were deemed significantly upregulated if the average natural logarithm fold change (logFC) exceeded 0.58 with an adjusted p-value less than 0.05. Conversely, genes exhibiting a logFC less than −0.58 with an adjusted p-value below 0.05 were classified as significantly downregulated.

### Gene Ontology (GO) Enrichment Analysis

To elucidate the biological functions of the identified DEGs, a GO term enrichment analysis was conducted using the clusterProfiler R package^36^. The functional annotations and subsequent visualizations were specifically confined to the GO Biological Process (GOBP) categories.

### Cell-Cell Communication Analysis

To deduce and examine intercellular communication networks, ligand-receptor interaction analysis was conducted utilizing the CellChat R package^37^. Overexpressed ligands and receptors identified within distinct cell populations were mapped onto a protein-protein interaction (PPI) network. CellChat extrapolates biologically significant cell-cell communications by assigning probability values to each interaction and conducting rigorous permutation tests. The resultant cellular communication networks and specific signaling pathways were visualized employing conventional circle and bubble plots.

### RNA Extraction and Quantitative Real-Time PCR (qPCR)

To corroborate the transcriptomic results, total RNA was isolated from patient and control PBMCs utilizing the TRIzol reagent in accordance with the manufacturer’s instructions. The extracted RNA was subsequently reverse-transcribed into cDNA. Quantitative real-time polymerase chain reaction (qRT-PCR) was conducted using the SYBR Green master mix on an ABI 7500 Real-Time PCR system. The relative expression levels of mRNA for the target genes (IFNG, IFNB1, MX1, and ISG15) were quantified employing the 2-ΔΔCt method, with GAPDH or ACTB used as the internal reference gene. Owing to limitations in RNA yield, qPCR validation was carried out on a representative subset of the initial cohort.

To validate the transcriptomic findings, total RNA was extracted from patient and control PBMCs using TRIzol reagent according to the manufacturer’s protocol. The RNA was reverse-transcribed into cDNA. qPCR was performed using SYBR Green master mix on an ABI 7500 Real-Time PCR system. The relative mRNA expression levels of target genes (IFNG, IFNB1, MX1, and ISG15) were calculated using the 2-ΔΔCt method, with GAPDH or ACTB serving as the internal reference gene. Due to available RNA yield limits, qPCR validation was performed on a representative subset of the original cohort.

### Enzyme-Linked Immunosorbent Assay (ELISA) Validation

ELISA was utilized to quantify the concentrations of IFN-γ, IFN-β1, MX1, and ISG15 proteins in plasma samples. The frozen plasma samples underwent centrifugation, and the assays were conducted in accordance with the manufacturers’ standard protocols. Absorbance readings were taken at 450 nm using a microplate reader. Standard curves were constructed employing a four-parameter logistic regression model to determine the concentrations of the target proteins, with all samples and standards analyzed in duplicate. Owing to the limited plasma volumes available from the acute cohort, ELISA measurements for certain targets, such as ISG15, were performed on representative subsets of the study population, as indicated in the corresponding figure legends.

### Statistics and Code Availability

R was used to perform the statistical analysis and visualizations. Wilcoxon rank-sum test or Student’s t-test was used to assess statistical significance. The following symbols represent significance: ns: *P* > 0.05; *: *P* ≤ 0.05; **: *P* ≤ 0.01; ***: *P* ≤ 0.001; ****: *P* ≤ 0.0001.

---

**Figure S1.**
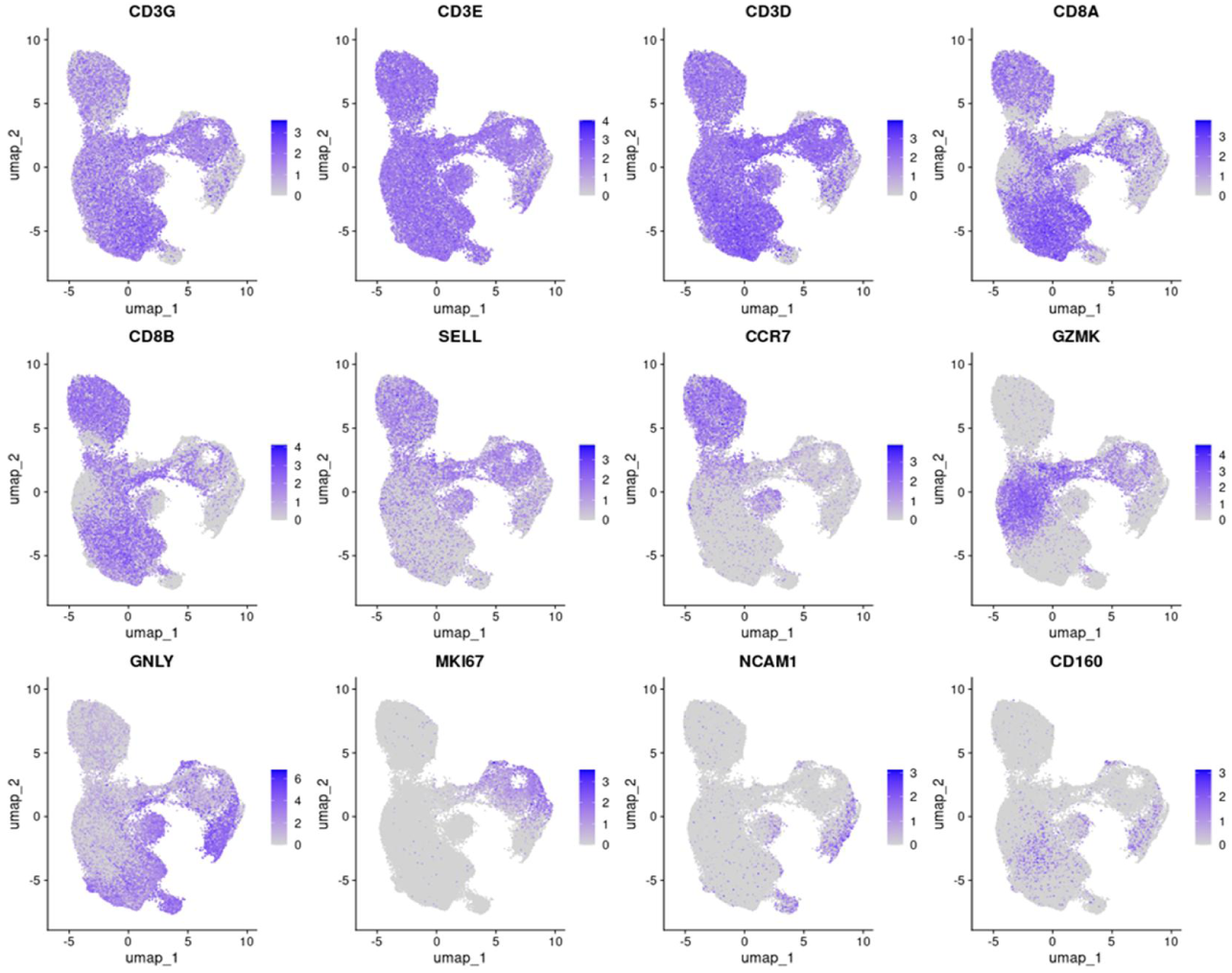
Marker for CD8^+^ T cell subsets.

**Figure S2.**
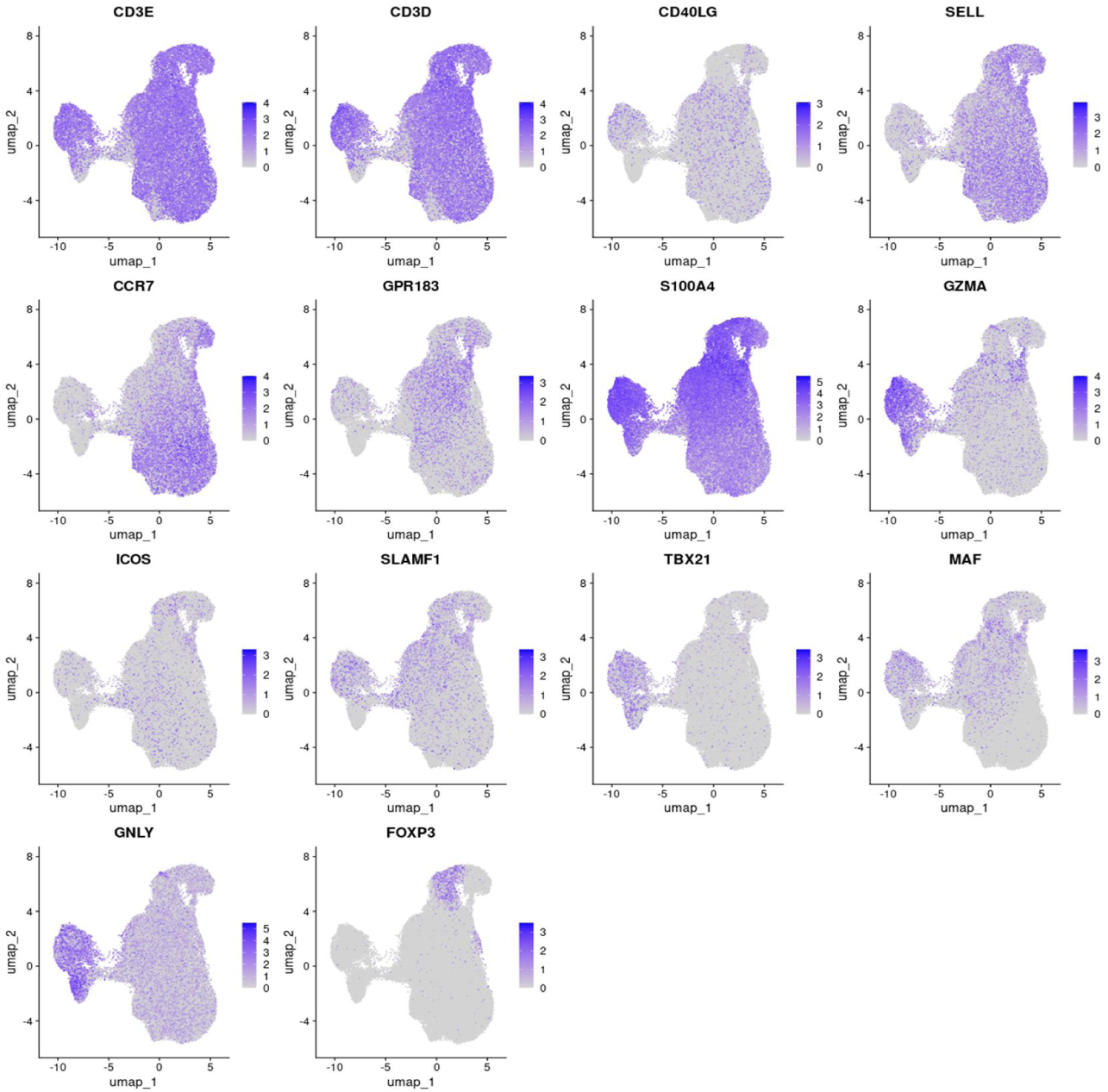
Marker for CD4^+^ T cell subsets.

## Notes

### Competing Interest Statement

The authors have declared no competing interest.

